# DMSP modulates virus-host interactions in the coral environment

**DOI:** 10.64898/2026.11.16.766974

**Authors:** Esther Rubio-Portillo, Rodrigo Sanchez-Martinez, Alejandro Díez-Martínez, Cristina López, Eva Mayol, Susanna Manrubia, Michael Steinke, Pedro J. de Pablo, Fernando Santos, Josefa Antón

## Abstract

Dimethylsulfoniopropionate (DMSP) is a ubiquitous organosulfur metabolite that plays central roles in marine ecosystems. It affects nutrient cycling, climate-relevant biogeochemical and microbial processes, including bacterial and viral infections of algal hosts, but its effect on bacteriophages has not been explored. In corals, DMSP is released under heat stress and attracts opportunistic *Vibrio* pathogens. Here we show that DMSP reshapes the infection dynamics of a virulent phage infecting the coral pathogen *Vibrio mediterranei*. DMSP initially decreased the number of infective viral particles through particle aggregation, extended the latent period and reduced burst size. However, prolonged infections ultimately yielded greater phage production, likely through DMSP-induced changes in the *Vibrio* envelope proteome. Consistent with these laboratory observations, DMSP also enhanced the proliferation of phage infecting *V. mediterranei* within the mucus of experimentally infected corals. Together, our findings suggest that DMSP released by stressed corals not only attracts bacterial pathogens but also promotes the amplification of their viral predators, revealing that host-derived metabolites can shape phage–pathogen interactions within animal microbiomes.

## INTRODUCTION

Dimethylsulfoniopropionate (DMSP) is an organosulfur compound produced in large amounts in the marine environment, with at least 8 billion tonnes estimated to be produced annually, mainly by phytoplankton as well as macroalgae, heterotrophic bacteria, and corals (Hopkins et al., 2023). It is a precursor of the climate-active gas dimethylsulfide (DMS), responsible for the typical smell of the sea, and exerts important biological effects. In addition to its well-known role as an osmolyte, DMSP can act as an antioxidant, cryoprotectant, chemoattractant, and as a crucial chemical signaling molecule, among other functions (Stefels et al., 2007). Besides its direct roles in DMSP-producing organisms (Wolfe & Steinke, 1996), once released into the environment by, for instance, viral lysis, DMSP can also serve as a source of reduced carbon and sulfur to some marine microbes. In addition, DMSP mediates biological interactions among different components of the complex systems, such as the chemosphere of the bloom-forming *Gephyrocapsa* (formerly *Emiliania*) *huxleyi* and the coral holobiont (Bratbak et al., 1995; Raina et al., 2010a).

DMSP released into the chemosphere of *G. huxleyi* regulates the outcome of the interactions between the algal host and its symbiotic bacteria since the exogenous application of DMSP increases *G. huxley*i susceptibility to bacterial infection (Barak-Gavish et al., 2018). This is the case for the algal associated *Sulfitobacter* D7, which can act both as a commensal or as an algicide pathogen, depending on the concentration of DMSP. DMSP also modulates the susceptibility of *G. huxleyi* to viral infection. Studies have shown that DMS and acrylate, the bacterial cleavage products of DMSP (released into the medium following viral infection), affect the infectivity of progeny viruses (Evans et al., 2006), ultimately reducing their infective titers and therefore decreasing the probability of infection of further cells.

Coral holobionts constitute another complex system in which DMSP is also prevalent. The coral holobiont comprises multipartite associations among the cnidarian host, its endosymbiotic dinoflagellate algae from the family Symbiodiniaceae, a diverse array of archaea and bacteria, viruses, and eukaryotes (fungi and non-Symbiodiniaceae protists) (Rohwer et al., 2002). The invertebrate host is one of the few animals able to synthesise DMSP, with concentrations increasing under stress conditions (up to 62 µM). DMSP is part of the coral chemosphere, which comprises a wide suite of metabolites identified as chemoattractants, antibacterial compounds and infochemicals (Ochsenkühn et al., 2018; Peixoto et al., 2017), thereby influencing microbial community structure and coral health status. For instance, despite its inability to use DMSP as a substrate, bacterial pathogens of the genus *Vibrio* may use this compound as a disease cue to target heat-stressed corals with increased susceptibility to infection (Garren et al., 2014).

Thus, DMSP can modulate the infectivity of both bacterial and viral pathogens toward their eukaryotic host. This raises the question of whether it can also mediate in a different type of pathogen–host interaction, namely that between bacteriophages (phages) and their bacterial hosts. To address this question, we chose the tripartite system constituted by the bacterial coral pathogen *Vibrio mediterranei*, its strictly virulent phage *Planavibriovirus adelos*, and its coral host, the invasive *Oculina patagonica*.

Heat stress has been shown to alter the microbial community of *O. patagonica*, particularly by increasing the abundance of *Vibrio* species (Martin-Cuadrado et al., 2024). Among these, *V. mediterranei* has emerged as one of the major threats, as it can induce coral bleaching. The strain used in this work was originally isolated in 2011 from a bleached colony of *O. patagonica* (Rubio- Portillo et al., 2014), while the phage *P. adelos* was isolated from the coral mucus layer following exposure to *V. mediterranei* (Rubio-Portillo et al., 2024). These phages are typically low- abundance specialists within the coral mucus microbiome and rapidly proliferate in the presence of their bacterial hosts, suggesting they may play an important role in coral defense against pathogenic infection (Rubio-Portillo et al., 2024). In this work, we unveil a new dimension of DMSP mediating phage–host dynamics both in pure culture and in the coral environment. We provide a mechanistic explanation on how DMSP affects virus-host interactions through a tradeoff between two contrary effects: the aggregation of viral particles and the increase of putative phage receptors in the host upon incubation with DMSP. This compound also strongly impacts corals exposed to the combined effects of the bacterial pathogen and its phage, both through the effect on the holobiont microbiome and on viral production. The discovery of this new function of DMSP provides support for the often-overlooked link between abiotic factors and virus-host interactions.

## RESULTS

### DMSP affects phage infectivity due to viral particle aggregation

The first clue pointing to the effect of DMSP on phage–bacteria interactions came from the observation that the incubation of virions of the phage *P. adelos* with different concentrations of DMSP impacted the formation of plaques, in a non-linear concentration-dependent manner (Fig. 1). As shown in Fig. 1b, incubation with 0.01-0.1 mM DMSP yielded a decrease of around 30% of PFU/mL compared to untreated viral-like particles (VLP). We hypothesized that this decrease was due to the aggregation of viral particles caused by DMSP, that would result in a decrease of the effective number of plaque-forming units (since one viral aggregate would yield only one plaque on a bacterial lawn). This hypothesis was supported by atomic-force microscopy (AFM) imaging of control and DMSP-treated VLPs, enabling a direct comparison between dispersed particles (Fig. 2a) and aggregated populations in the presence of DMSP (Fig. 2b).

**Fig. 1:**
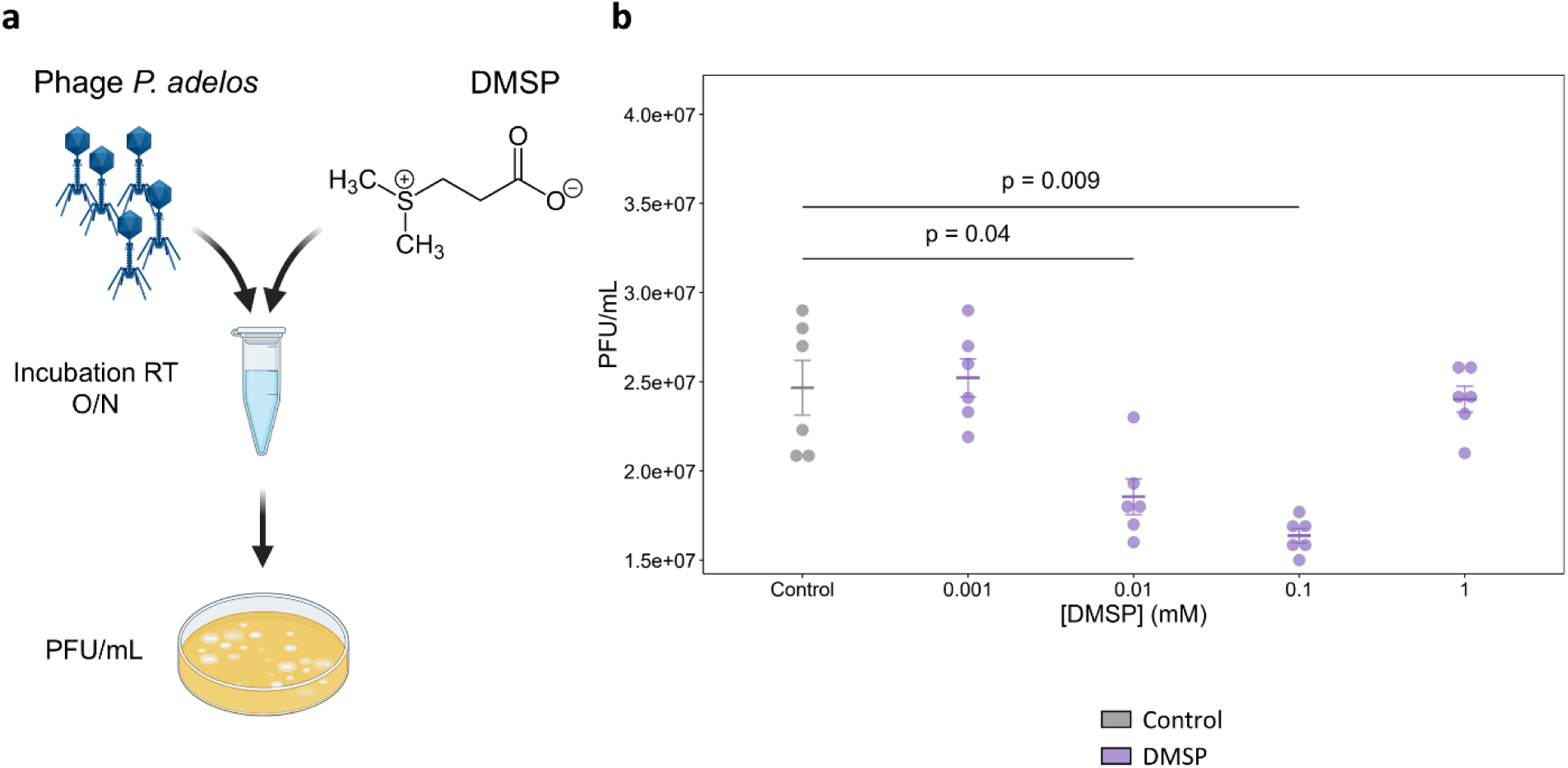
DMSP decreases phage infectivity in a concentration-dependent manner. a, Schematic representation of the experimental design used to assess the effect of DMSP on phage infectivity. Phage particles were incubated with the indicated DMSP concentrations before infective particles were quantified by plaque assay using *V. mediterranei* as the host. **b,** Infective phage titres, expressed as plaque-forming units PFU/ml, after incubation with 0, 0.001, 0.01, 0.1 or 1 mM DMSP. Dots represent individual biological replicates; horizontal lines and error bars indicate the mean ± SEM. For each phage, every DMSP concentration was compared with the untreated control using two-sided unpaired Welch’s t-tests, followed by Bonferroni correction for the four comparisons. Only statistically significant adjusted P values are shown; P < 0.05 was considered significant.

**Fig. 2:**
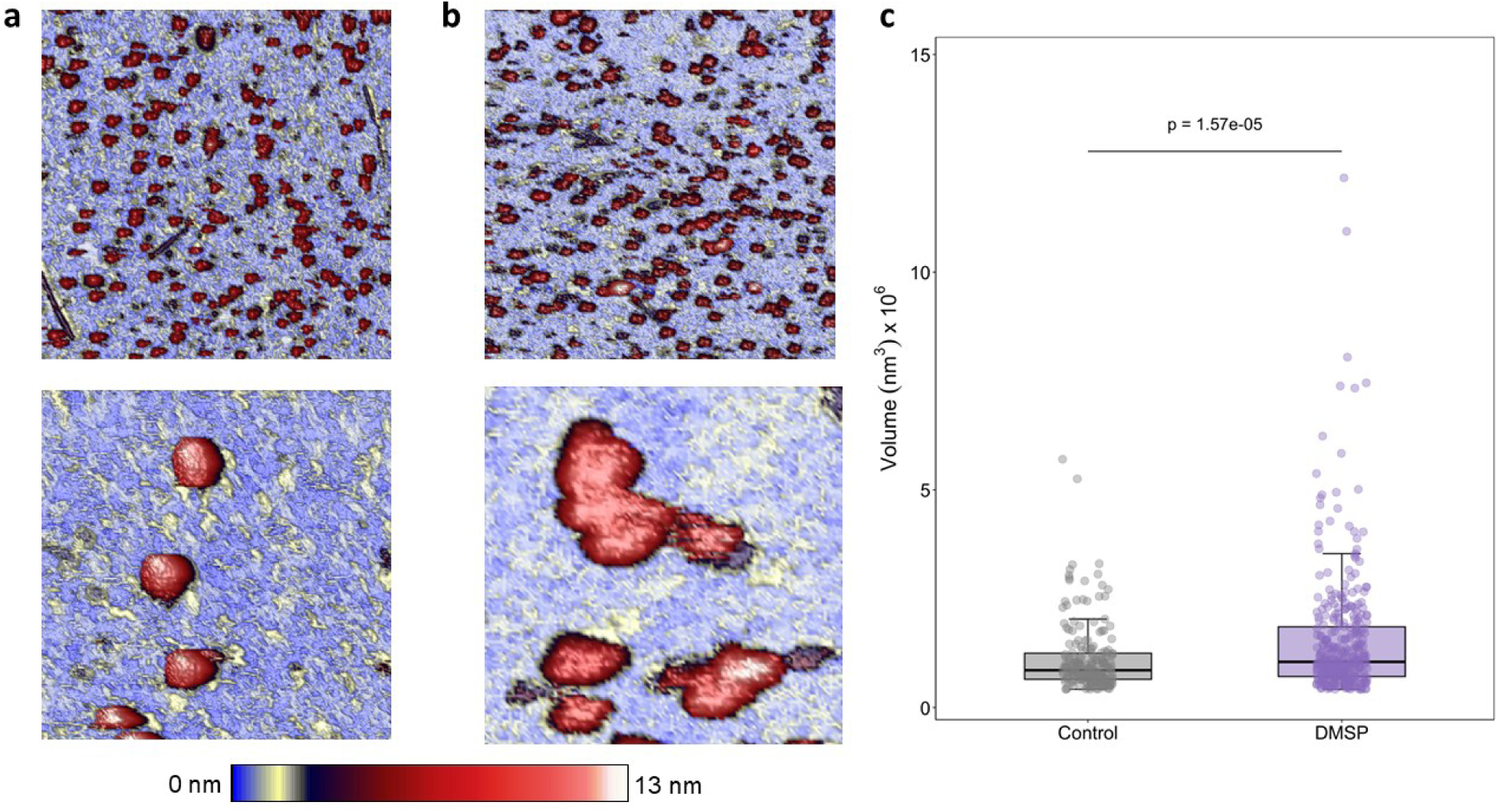
DMSP promotes aggregation of *P. adelos* viral particles. a,b, Representative atomic- force microscopy (AFM) images of untreated phage particles **(a)** and phage particles incubated with 0.1 mM DMSP **(b)**. Upper and lower images show 4 × 4 μm² and 1 × 1 μm² fields of view, respectively. **c,** Distribution of the apparent volumes of particles detected in control and DMSP- treated samples. Dots represent individual particles; boxes indicate the median and interquartile range, and whiskers extend to 1.5 × the interquartile range. Groups were compared using a two- sided unpaired Welch’s t-test. The exact P value is shown; P < 0.05 was considered significant.

This qualitative observation was supported by quantitative AFM image analysis (Fig. 2c), which revealed a significant increase in the apparent volume of detected viral particles following DMSP treatment (*t-test*, p <0.001). Particle volume distributions were used as a first-order approach to identify aggregation, as larger apparent volumes corresponded to multiple virions detected as a single object by the analysis software. Because individual virions and aggregates can exhibit substantial morphological variability in AFM images, aggregates were identified using a volume- based threshold rather than visual morphology alone. Thus, particles falling above the upper whisker of the control distribution, corresponding to values outside the 1.5 × IQR range, were classified as aggregates. Based on this criterion, the control sample contained approximately 10% aggregates, whereas the DMSP-treated sample reached ∼25%. Importantly, the volume distribution of the DMSP-treated sample is globally shifted towards higher values, indicating not only a higher fraction of aggregates but also larger aggregate sizes. This suggests that many of these structures were composed of more than two virions, implying that the extent of aggregation is underestimated when considering only aggregate counts. Consequently, DMSP-induced aggregation was likely more pronounced than the ∼25% value derived from the threshold-based classification.

### DMSP induces a reduction of burst size followed by higher viral production

One-step infection curves, provide three relevant parameters about the phage life cycle: number of non-attached VLPs at the beginning of the infection experiment (i.e. free VLPs that remain infective after treatment), the latent period (i.e. time from the attachment until cell lysis) and burst size (average number of virions produced by infected cell). When viral particles of phage *P. adelos* were incubated in DMSP prior to infection, an effect could be observed in either of these aspects. First, free viral particles were around 4 times higher in untreated control than when viruses were treated with DMSP, which is not unexpected given the results from AFM, since an aggregate of virions forms only one individual plaque. DMSP also induced a delay in the latent period and a decrease in burst size (calculated as in Clokie & Kropinski, 2009) from 455 to 356 virions/cell (Fig. 3a), likely due to a decrease in the effective number of virions per cell or multiplicity of infection (MOI). This suggests that DMSP decreased the effective “virulence” of phage *P. adelos* against *V. mediterranei*. These effects were also observed when DMSP treatment was applied to the cells (instead to phages) prior to the infection, as discussed below.

**Fig. 3:**
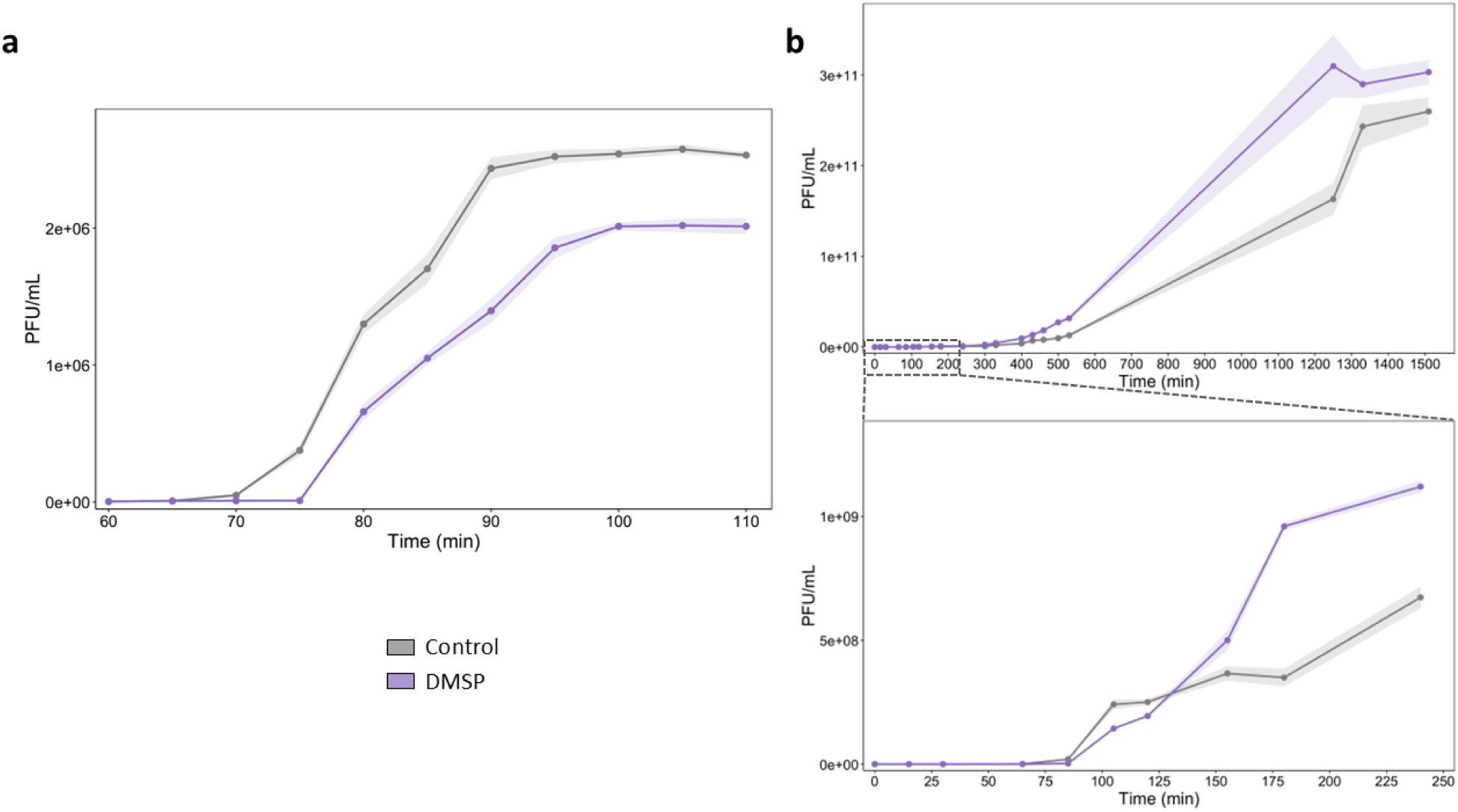
DMSP delays the initial replication cycle of *P. adelos* but enhances viral production during prolonged infection. a, One-step growth curves of *P. adelos* infecting *V. mediterranei*. Phage particles were preincubated for 12 h in the absence (control) or presence of 0.1 mM DMSP before infection at a multiplicity of infection of 0.1. Infective phage titres were quantified by plaque assay over 110 min and are expressed as plaque-forming units PFU/ml. **b,** Extended infection kinetics under the same conditions, showing viral production during prolonged incubation. Points represent the mean of three independent biological replicates; shaded areas indicate the mean ± SEM.

However, when the cultures initially used to assess the one-step growth curve were incubated beyond the 110 min of the one-step curve experiment, a counter-intuitive result was observed: in infection experiments treated with DMSP, a smaller burst size (455 PFU/cell for the control and 346 PFU/cell for the DMSP-treated) at the beginning was accompanied with an increased production of viruses after 150 min incubation (2.6x10^11^ PFU/mL for the control and 3x10^11^ PFU/mL for the DMSP-treated) (Fig. 3b). Modelling demonstrated analytically that neither the changes in burst size and latent period, nor the bacterial growth occurring during the extended latent period, are sufficient to produce this reversal — the latter would require an unrealistically fast doubling time (<12 min), well below the measured value (37 min) — implying that DMSP must be affecting some host processes or the interaction with the virus, perhaps enhancing the efficiency of infection itself (Supplementary Material). This implied that there is either an effect of DMSP on the bacterial host growth and/or on the infection rate beyond the direct effect on the viral particles, that had not been considered given de absence in the *V. mediterranei* genome of genes involved in the use of DMSP as a nutrient. In fact, the only DMSP-related gene detected in the *V. mediterranei* genome coded for the DMSP lyase DddP. This protein can cleave DMSP, releasing DMS, although with a very low activity, which suggests that DMSP is not its native substrate (Interpro entry: IPR050020).

Monitoring growth of *V. mediterranei* cultures supplemented with 0.1 mM DMSP indicated that this compound exerted a minor effect on *V. mediterranei* growth in the first 4h. However, these differences lacked statistical significance (1.5x10^9^ CFU/mL for the control and 1.7x10^9^ CFU/mL for the DMSP-treated) (Fig. 4a). A slight effect was perceptible after 150 min after the addition of DMSP, a time period exceeding the time required to detect the first synchronous burst of cells in the one-step curve of the infected culture, that was around 75 min (Fig. 3a). The effect of DMSP on *V. mediterranei* growth was then explored through proteomics.

**Fig. 4:**
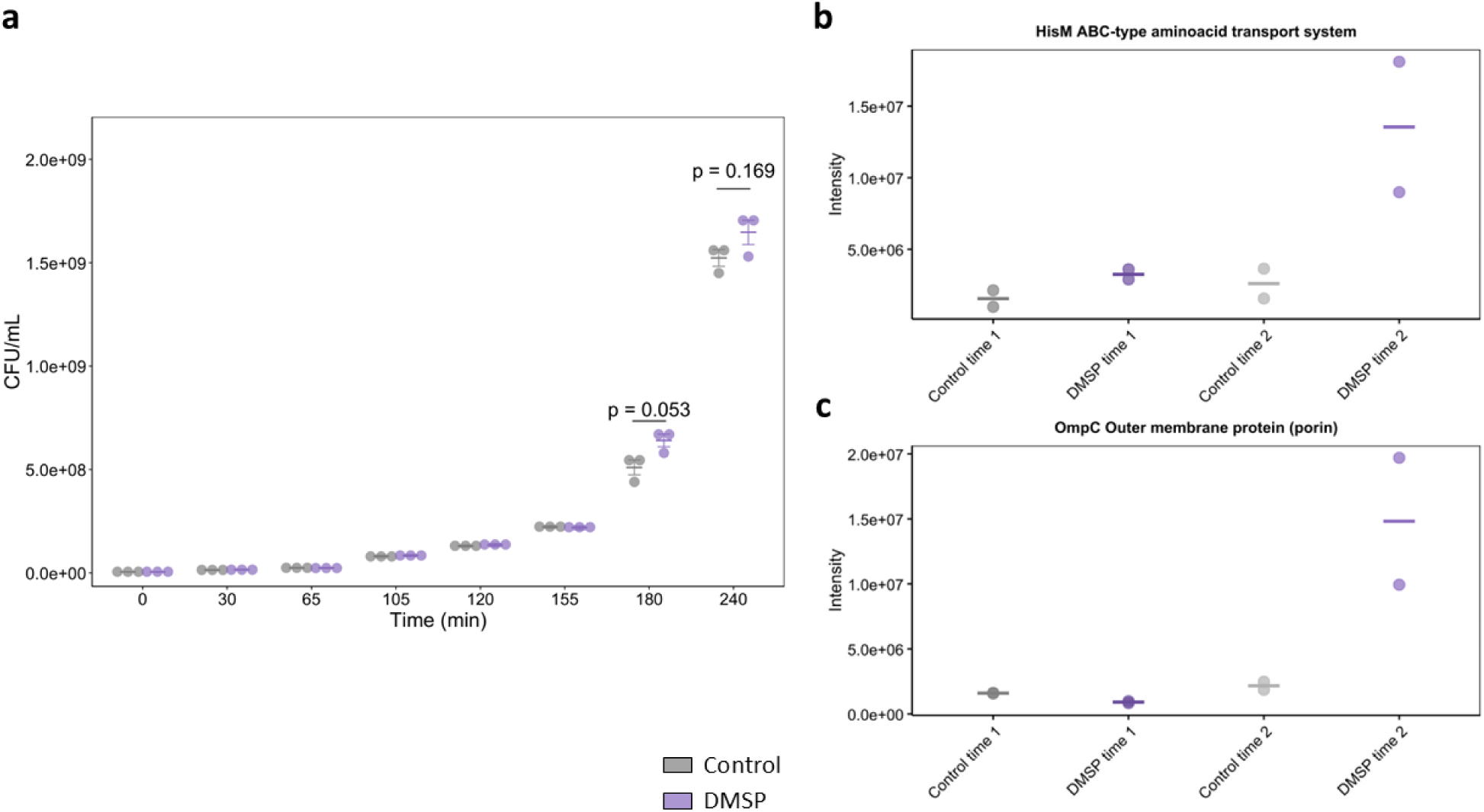
DMSP alters the envelope proteome of *V. mediterranei* with only a minor effect on bacterial growth. a, Growth of *V. mediterranei* cultures in the absence (control) or presence of 0.1 mM DMSP, monitored as colony-forming units CFU/ml over time. Each point represents one of three independent biological replicates; horizontal lines and error bars indicate the mean ± SEM. **b,c,** Comparative analysis of the *V. mediterranei* envelope proteome after incubation in the absence or presence of 0.1 mM DMSP, highlighting differentially abundant proteins, including a HisM ABC-type amino acid transport protein and an OmpC porin. Horizontal lines represent the mean of two independent biological replicates. Samples were taken at Time 1 and Time 2 which correspond to 2h and 3h after inoculation, respectively.

### Exposure to DMSP increases viral production due to changes in the host’s envelope proteome

A proteomic study of the envelope proteins expressed at the beginning of the exponential phase at 2 and 3 h post inoculation with and without added DMSP was carried out. Results (Fig. 4b, c) indicated that there was an increase in the abundance of at least two proteins that could be involved as phage receptors: a HisM ABC-type aminoacid transport protein and a porine of the OmpC family. Porin OmpF is known to be involved in phage-host recognition in *Vibrio alginolyticus* (Skliros et al., 2021), while OmpC itself also mediates phage-host recognition in *Escherichia coli* and other members of the *Pseudomonadota* (Ho & Slauch, 2001; Terasaki et al., 2026; Washizaki et al., 2016). Notably, the increase in infection efficiency required by the population-dynamics model to reproduce the observed ∼2-fold reversal, closely matched the fold-change threshold used to call these proteins significantly regulated (Supplementary Material).

This indicated that the effect of DMSP on the interaction between *V. mediterranei* and its phage detected in the infection experiments (Fig. 3) could be due to the modification of envelope receptors since they showed increased production when the host was in contact with this compound (Fig. 4). Thus, there was a tradeoff between two contrary phenomena: the direct (on aggregation) and indirect (through modulation of host physiology) effects of DMSP on the outcome of the interaction between *V. mediterranei* and its phage.

### DMSP modulates viral production and the coral microbiome, increasing the relative abundance of ‘ultrasmall’ prokaryotes

In order to assess whether DMSP also had an effect on parts of the coral holobiont due to the interaction of *V. mediterranei* and its phage in their natural context (Garren et al., 2014; Raina et al., 2010a), 36 pieces of corals separated into replicates of six each per aquarium tank were submitted to different treatments of combinations of *Vibrio*, phage and DMSP (Fig. 5a). After 17 days of incubation, coral mucus was obtained and used to quantify viral particles and to monitor changes in the microbial communities using metabarcoding. Overall, although the treatment replicates displayed very high heterogeneity in chlorophyl *a* (Chl *a*) and Shannon entropy (Extended Data Fig. 1), two clear observations with statistical support could be derived from the experiment regarding the effects on viral abundance and microbiome composition.

**Fig. 5:**
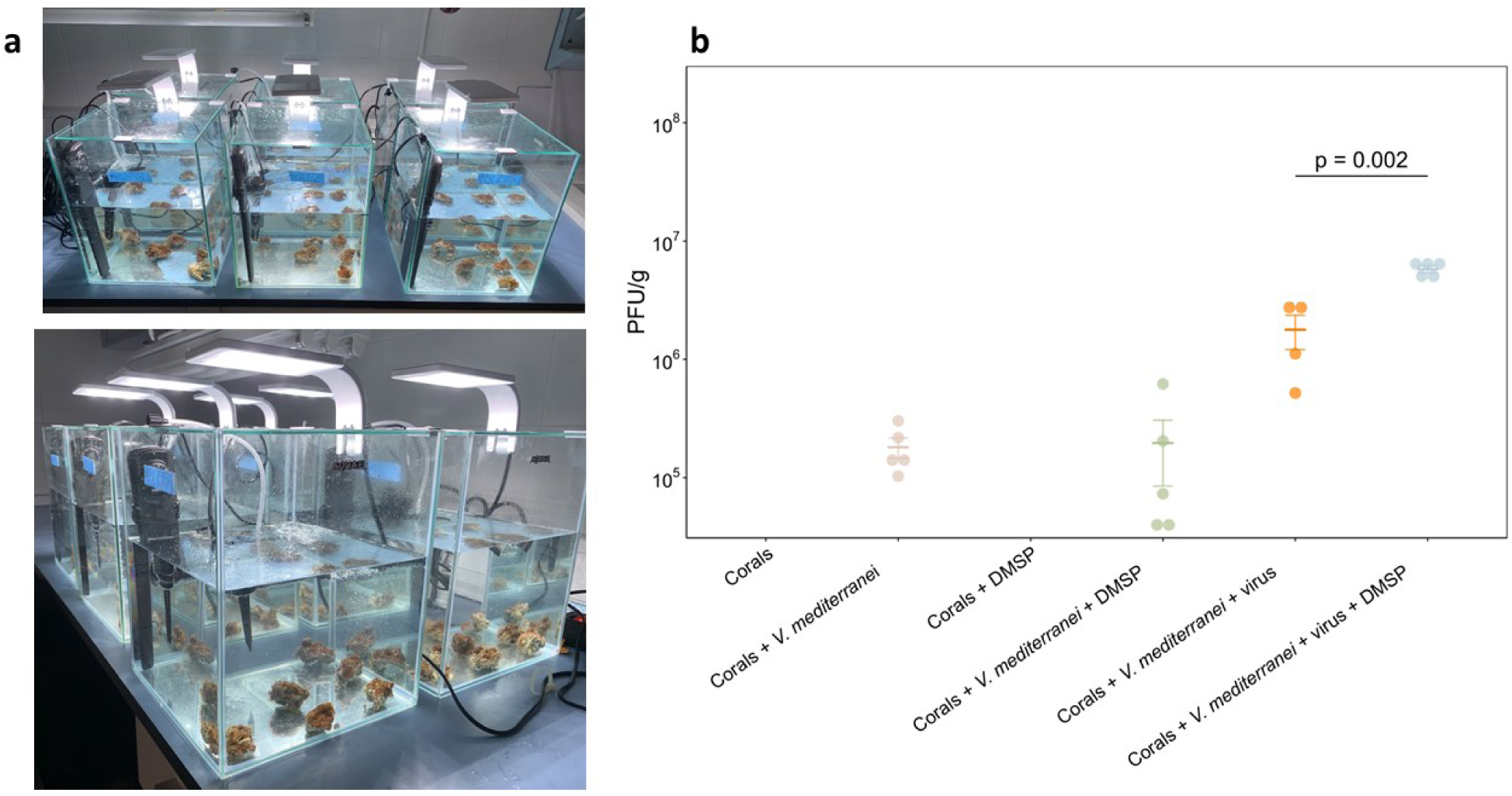
DMSP enhances the proliferation of phages infecting *V. mediterranei* in coral mucus. a, Fragments of *O. patagonica* were assigned to six treatment aquaria: no addition (control), *V. mediterranei* alone, DMSP alone, *V. mediterranei* with DMSP, *V. mediterranei* with *P. adelos*, and *V. mediterranei* with *P. adelos* and DMSP. DMSP was added at 0.1 mM, and bacterial and phage inocula were added at final concentrations of 10³ CFU/ml and 10² PFU/ml, respectively. Each aquarium contained six coral fragments, and samples were collected after 17 days. **b,** Abundance of phages infecting *V. mediterranei* in coral mucus at the end of the experiment, expressed as PFU/g. Dots represent individual coral fragments; horizontal lines and error bars indicate the mean ± SEM. The two treatments containing both *V. mediterranei* and *P. adelos*, with or without DMSP, were compared using a two-sided unpaired Welch’s t-test. The exact P value is shown; P < 0.05 was considered significant.

When coral treated with *Vibrio* and phage were supplemented with DMSP, phages able to infect *V. mediterranei* increased consistently in the coral mucus, as determined by the number of PFU/g (Fig. 5b). Hence, DMSP exerted the same effect on the interaction of *V. mediterranei* with its phage as what was observed in culture, namely an increase of viral production. Two additional observations can be derived from data in Fig. 5: phages infecting *V. mediterranei* were naturally present in the coral mucus, and the PFU added to the aquaria could thrive in the mucus compartment.

The impact of the different treatments on microbial community was ascertained by 16S rRNA gene metabarcoding. PERMANOVA indicated that there was not a single factor whose effect on the microbial community could be statistically supported. Instead, there was a significant effect of the interaction between the factors phage and DMSP. The effect of this interaction on the mucus microbiome was further investigated by Compositional Data Analysis (CoDA) using the Logratio and Glmnet packages, which assess changes in the relative composition of the microbiome and identify the Operational Taxonomic Units (OTUs) driving the observed separation between treatments. These analysis detected a total of 70 OTUs responsible for the separation of corals infected with *V. mediterranei* and its phage depending on whether they had been treated with DMSP or not (Fig. 6). Most of these OTUs belonged to the so-called rare biosphere (below 0.1% relative abundance (Pedrós-Alió, 2012)), and did not feature genera typically considered as DMSP-metabolizing bacteria, such as *Roseobacter*. Among the 21 OTUs which increased their relative abundance due to DMSP exposure, 11 corresponded to ultrasmall prokaryotes, belonging either to the Patescibacteria or the Nanoarcheota, while only 1 OTU of these phyla was detected among the 36 phylotypes that decreased in aquaria with DMSP. Overall Patescibacteria and Nanoarchaeota increased ten-fold from 0.87 to 9.17% relative abundance, with some OTUs entering the abundant biosphere upon DMSP treatment.

**Fig. 6:**
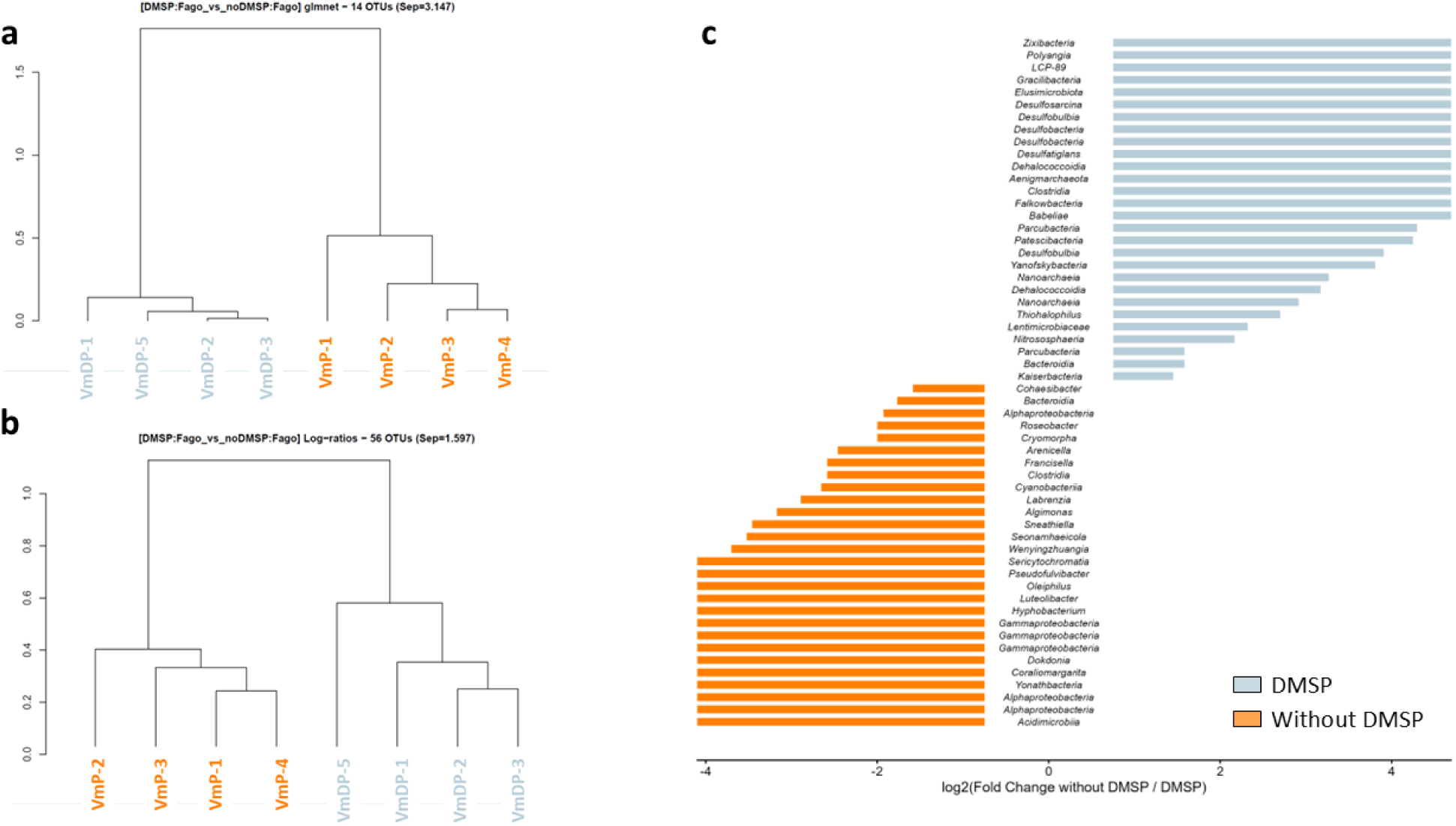
DMSP alters the relative abundance of coral mucus-associated OTUs in the presence of *V. mediterranei* and its phage. a, Hierarchical clustering of coral mucus samples based on the relative abundance profiles of 14 OTUs selected using glmnet and **b,** 56 OTUs selected using the log-ratio approach. All samples were incubated with *V. mediterranei* and its phage. In both dendrograms, samples treated with DMSP (VmDP, grey) formed a distinct cluster from samples incubated without DMSP (VmP, orange). **c,** Log₂ fold changes in OTU relative abundance between treatments without and with DMSP. Each bar represents an OTU selected using the log-ratio approach and is labelled with its taxonomic assignment. OTUs with zero relative abundance in any sample included in the comparison were excluded from the fold-change calculation.

When the interaction between phage and DMSP was explored by analyzing the OTUs responsible for the differences due to phage treatments, Logratio detected 29 OTUs responsible for the differences (Fig. 7) that, again, mostly corresponded to components of the rare biosphere. Additionally, the presence of phage was associated with a seven-fold increase in ultrasmall prokaryotes, from 0.4 to 2.6% relative abundance.

**Fig. 7:**
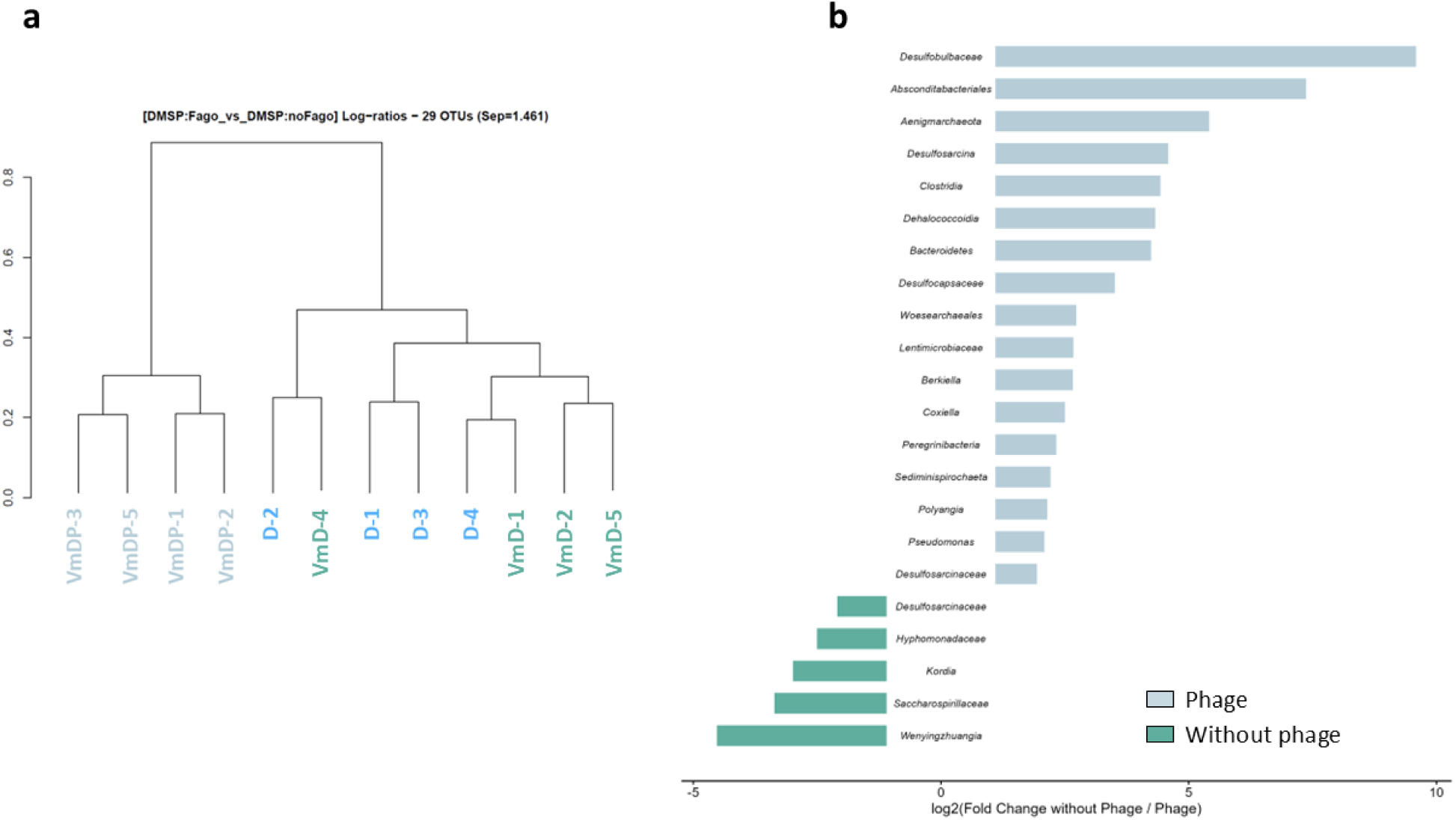
Phage exposure alters the relative abundance of coral mucus-associated OTUs in the presence of DMSP. a, Hierarchical clustering of DMSP-treated coral mucus samples based on the relative abundance profiles of 29 OTUs selected using the log-ratio approach. Samples exposed to *V. mediterranei* and its phage (VmDP, grey) formed a distinct cluster from samples treated with DMSP alone (D, blue) or with DMSP and *V. mediterranei* (VmD, green). **b,** Log₂ fold changes in OTU relative abundance between treatments without and with added phage under DMSP supplementation. Each bar represents an OTU and is labelled with its taxonomic assignment. OTUs with zero relative abundance in any sample included in the comparison were excluded from the fold-change calculation, leaving 22 of the 29 selected OTUs in panel **b**.

## DISCUSSION

This study unveils a previously unrecognized role of DMSP as a modulator of phage–bacteria interactions, which adds to the list of functions that this metabolite has in the marine environment.

Classical studies on the interaction of the coccolithophore *G. huxleyi* with its viruses showed that the DMSP cleavage products, including DMS and acrylate, could act as a chemical defense against viral infection (Evans et al., 2006; Schroeder et al., 2002). Here, we show that DMSP itself may modulate phage–bacteria interactions through a combination of a direct physical effect on the viral particles and indirect effects mediated by changes in host physiology, through a trade-off between two opposite outcomes.

DMSP causes virions to aggregate, shifting the particle-size distribution toward larger structures, as shown by AFM. This aggregation likely explains the initial reduction in phage titers due to the reduction of apparent infective units, delaying latent periods and reducing the burst size in one- step growth curves. This could be considered an initial antiviral effect of DMSP, mirroring that observed in the *G. huxleyi*- virus system, but operating through a completely different mechanism. However, as the infection proceeds, there is an increase in viral production after 110 min, suggesting that DMSP also affects the bacterial host rather than exclusively the phage, a conclusion also reached independently from the dynamical model (Supplementary Material). Incubation with DMSP alters the envelope proteome of *V. mediterranei*, increasing the abundance of proteins that likely act as phage receptors, including an OmpC-family porin, a class of proteins frequently involved in phage adsorption (Ho & Slauch, 2001; Terasaki et al., 2026; Washizaki et al., 2016) and other envelope proteins including a putative HisM ABC-type aminoacid transport protein. Thus, *V. mediterranei* responds to DMSP despite lacking canonical DMSP-catabolic pathways. These findings suggest that this compound is not acting as a nutrient but as a physiological signal, modulating the bacterial growth and its susceptibility to infection.

This interpretation is in good agreement with the well-established role of DMSP as an infochemical for coral-pathogenic *Vibrio* species. Quorum sensing autoinducer activity in a strain of *V. shilonii* (the former name of *V. mediterranei*) isolated from bleached corals increased in response to 0.005 mM DMSP but decreased in response to 0.05 mM (Li et al., 2017) . Likewise, the coral pathogen *V. coralliilyticus* uses DMSP as a chemotactic cue to locate heat-stressed corals, despite being unable to metabolize it (Garren et al., 2014). In this system, DMSP concentrations increased fivefold under thermal stress, and bacterial attraction scaled nonlinearly with DMSP levels.

DMSP is known to mediate virulence in algicidal members of *Roseobacteraceae* such as *Sulfitobacter* D7, where it increases the expression of transporters and virulence-related genes, enhancing the pathogenicity toward *G. huxleyi* (Barak-Gavish et al., 2018). Despite the biological differences among the *Sulfitobacter*–*G. huxleyi*, *Vibrio*–phage, and *Vibrio*–coral systems, they all point to a common role of DMSP as a ‘behavioral’ signal that mediates parasite–prey interactions, even in organisms unable to catabolize it.

The aquarium experiment demonstrated the complexity of the effects of DMSP on the *Vibrio*– phage system within its natural environment, the coral holobiont. Indeed, the effects of DMSP on the coral microbiome were detected primarily in combination with phage activity, as indicated by a significant phage×DMSP interaction in the PERMANOVA analysis, whereas no significant effects of the single factors were detected. This reinforces the idea that chemical cues and viral dynamics are tightly connected in coral ecosystems. The microbiome analyses revealed that the effect of DMSP was not restricted to *Vibrio* or other classical DMSP-metabolizing taxa. Instead, the combined effects of phage infection and DMSP reshaped the rare biosphere, including ultrasmall prokaryotes. These microbes have been detected as part of the rare biosphere of the coral (Campos et al., 2022). In the words of Campos et al., CPR and DPANN have an overlooked role in corals’ microbial community structure. Furthermore, our results are in good agreement with a recent meta-study of 2860 metagenomes spanning nine ecosystems that found that rare taxa, primarily CPR and DPANN, disproportionately shape microbial community dissimilarities across global ecosystems (Ferreira et al., 2025).

The abundance of viruses infecting *V. mediterranei* increased in the mucus of corals exposed to *Vibrio*, extending our previous observations from culture to an experimental infection model. In our previous study, we proposed that lytic phages targeting coral pathogens became more abundant in infected coral mucus and could constitute an important first line of defense against bacterial invasion (Rubio-Portillo et al., 2024). Here, we further show that phage proliferation is enhanced in the presence of DMSP. Given that DMSP production increases in stressed corals (Raina et al., 2013) and that this metabolite serves as a key chemical cue for pathogenic *Vibrio* spp. (Garren et al., 2014), these findings provide a plausible mechanistic explanation for the enrichment of pathogen-specific phages during the infection. This adds a new dimension to our previous hypothesis by suggesting that phage-mediated protection is not merely the result of a standing population of lytic viruses in the mucus, but rather a dynamic response that may be reinforced by chemical signals released by the coral itself. Consequently, stressed corals could enhance the abundance of phages targeting opportunistic pathogens precisely when they become most susceptible to infection, thereby strengthening an intrinsic phage-based defense mechanism within the coral holobiont.

Overall, our work demonstrates that DMSP-rich systems, such as coral mucus, emerge as environments where phage-bacteria interactions are actively modulated by infochemicals. This adds a new dimension to our understanding of microbial interactions, indicating that chemical and viral ecologies are deeply intertwined.

## METHODS

### Bacterial strains, viruses, growth conditions and DMSP treatments

*Vibrio mediterranei* strain Vic-Oc-097 (CECT 30098), originally isolated from a bleached colony of the coral *Oculina patagonica* (Rubio-Portillo et al., 2014), and the phage *Planavibriovirus adelos* strain MVO1_097.13, isolated from coral mucus (Rubio-Portillo et al., 2024), were used in this study. *V. mediterranei* was grown aerobically at 28 °C in Luria–Bertani (LB) medium supplemented with 3% (w/v) NaCl, with shaking at 120 rpm. Viral stocks were produced by infecting exponentially growing *V. mediterraei* cultures and incubating them for 12 h at 28 °C with shaking at 120 rpm. The lysate was subsequently centrifuged to remove bacterial debris, filtered through a sterile 0.22-µm pore-size membrane, and stored at 4°C until use. Phage titers were determined using the double-layer agar (“plaque assay”) method and expressed as plaque-forming units (PFU) mL⁻¹. For bacteriophage enumeration, 10-fold serial dilutions of coral mucus were mixed with a mid-exponential-phase bacterial culture and added to 4 mL of molten LB soft agar (0.7% agar) supplemented with 3% (w/v) NaCl. The mixture was poured onto LB supplemented with 3% (w/v) NaCl agar plates and incubated overnight at 30 °C.

To assess whether dimethylsulfoniopropionate (DMSP) affects phage infectivity, viral suspensions were incubated with different concentrations of DMSP (Sigma, Missouri, USA) (1, 0.1, 0.01, and 0.001 mM) at room temperature for 12 h. Following incubation, treated phages were mixed with mid-exponential-phase cultures of *V. mediterranei* and quantified using the double-layer agar plaque assay. Changes in plaque-forming units (PFU) were used as a proxy for phage infectivity. The concentration of DMSP was chosen according to the levels detected in the natural environment from which *V. mediterranei* and its phage were isolated (Raina et al., 2010b).

To evaluate whether DMSP influences the growth of *V. mediterranei*, triplicate cultures were grown at 28 °C in the presence or absence of 0.1 mM DMSP. Bacterial growth was monitored by plate counts of *Vibrio* onto thiosulfate citrate bile salts sucrose (TCBS) agar (Pronadisa, Spain).

### Atomic force microscopy (AFM)

The viral stock was maintained at 4 °C at a concentration of 10¹² particles/ml in SM buffer (50 mM Tris-HCl, 100 mM NaCl, 8 mM MgSO₄, 0.005% glycerol, pH 7.5). For the experiments, the sample was diluted 25-fold with the same buffer. For aggregation assays, DMSP was added to a final concentration of 0.1 mM followed by brief vortexing and incubation at 4 °C for 24 h. For AFM imaging, 30 µl of each sample was deposited onto freshly cleaved mica and allowed to adsorb for 20 min under ambient conditions. The surface was then gently rinsed three times with the corresponding buffer (either SM buffer or SM supplemented with 0.1% DMSP for aggregation conditions) to remove non-adhered viral particles. The liquid cell was subsequently filled to a final volume of 120 µL to ensure stable imaging under fully hydrated conditions. Medium-stiffness qp- BioAC cantilevers (rectangular geometry, nominal tip radius <10 nm) were used for all measurements. The spring constant (nominal value 0.09 N/m) was calibrated individually prior to each experiment using Sader’s method in air (Sader et al., 1999). Image acquisition and analysis were performed with WSxM software (Horcas et al., 2007). AFM measurements were carried out using a system from Nanotec Electrónica S.L. (Madrid, Spain), operating in an updated implementation of Jumping Mode Plus (Ortega-Esteban et al., 2012). In this approach, the probe is driven vertically towards the surface until a predefined normal force is reached. It is then retracted before lateral displacement to the next pixel, where the cycle is repeated. This point-by- point acquisition enables accurate force control while significantly reducing lateral forces, thereby preserving the structural integrity of the viral particles during imaging.

### One-step growth curve

Bacteriophage *P. adelos* was incubated at room temperature for 12 h in the absence of host bacteria, either with or without DMSP (0.1 mM). After incubation, burst size was determined after carrying out a one-step growth experiment as detailed in Clokie and Kropinski (2009), with slight modifications. Briefly, a culture of *V. mediterranei* at mid-exponential phase (OD₆₀₀ = 0.5, 3 hours) was infected with *P. adelos* (preincubated in parallel with and without DMSP as described above) at a multiplicity of infection (MOI) of 0.1 and incubated 5 min at 28 °C. The infections were diluted 10²-, 10³-, and 10⁴-fold in LB medium supplemented with 3% (w/v) NaCl, either in the presence or absence of 0.1 mM DMSP, and incubated at 28 °C. Aliquots of 150 μL were taken every 5 min for 110 min and the numbers of PFU were determined by plaque assay. At time 0 min, an additional 1 ml aliquot was collected, treated with a few drops of chloroform, and used to quantify non-adsorbed bacteriophages by plaque assay.

To assess the long-term effect of DMSP on viral production, a second experiment mirroring the one described above was performed, with incubation extended to 24 h and multiple aliquots collected for PFU quantification. All experiments were performed in triplicate.

### Proteomics

*V. mediterranei* cultures grown in the presence or absence of DMSP (0.1 mM) were used for envelope protein extraction. Cultures were harvested at two sampling points, 3 hours after inoculation when mid-exponential phase was reached (OD₆₀₀ = 0.5, T_0_, this is the time when DMSP was added in the burst-size experiment) and after 120 minutes (T_1_), by centrifugation at 4,000×g for 15 min at 4 °C. The cell pellets were washed three times with 40 ml of sterile saline solution (0.15 M NaCl) and subsequently resuspended in 5 ml of sterile seawater. Cells were disrupted by intermittent sonication (25 s pulses at 10% amplitude). Unbroken cells and cellular debris were removed by centrifugation at 5,000×g for 20 min at 4 °C, and the supernatant was further centrifuged at 100,000×g for 40 min at 4 °C to pellet membrane fractions. The resulting pellet was resuspended in 10 ml of 2% (w/v) sodium lauryl sarcosinate (Sigma, Missouri, USA) and incubated at room temperature for 1 h. The suspension was then centrifuged at 100,000×g for 40 min at 4 °C. The final pellet was resuspended in 200 µL of sterile seawater and stored at -20 °C until further analysis.

For the tryptic in-solution digestion, 30 micrograms of precipitated protein sample were dissolved in 25 μl of 6 M urea. Reduction was achieved with 5μl of 0.2 M dithiothreitol, followed by incubation for 1 h at 37 °C and S-alkylation with 20μl of 0.2 M iodoacetamide, followed by incubation for 1 h in the dark at room temperature. Before tryptic digestion, 25 mM ammonium bicarbonate buffer was added to decrease the concentration of urea to 0.6 M. For in-solution digestion, trypsin was added to the protein mixture at an enzyme-to-substrate ratio of 1:30 (w/w). After incubation at 37 °C for 16 h, additional trypsin (1:60, w/w) was added to the sample and incubation was continued for 5 h to ensure complete digestion. Tryptic peptides were dried down in a Speed-Vac benchtop centrifuge and resuspended in 5% acetonitrile and 0.5% trifluoroacetic acid. The resulting peptides were desalted with PepClean C-18 Spin Columns (Agilent Technologies, CA, USA) in batches of 30 μg of protein according to manufacturer recommendations. Eventually, eluted peptides were dried down in a Speed-Vac benchtop centrifuge and resuspended in 10 µL of first LC mobile phase (5% acetonitrile and 0.1% formic acid).

### LC-MS/MS analysis

Proteomic analyses were carried out in the Research Technical Services at the University of Alicante, by means of LC-MS/MS after tryptic digestion of the protein extract. To perform label-free relative quantitation, four technical replicates from each sample were analyzed. Peptide separation was performed using an Agilent 1290 Infinity LC system coupled to the 6550 Accurate-Mass QTOF (Agilent Technologies, CA, USA) with electrospray interface (Jet Stream Technology) operating in positive-ion mode (3,500 V) and in high-sensitivity mode. The best conditions for the electrospray interface were: gas temperature 250 °C, drying gas 14 L/min, nebulizer 35 psi, sheath-gas temperature 250 °C, sheath-gas flow 11 L/min. Samples were injected (10 μL) on an Agilent AdvanceBio Peptide mapping column (2.1 × 250 mm, 2.7 μm) (Agilent Technologies, CA, USA) with a 3 to 40% gradient of solvent B (0.1% formic acid in 90% acetonitrile) for 140 min operating at 50 °C and a flow rate of 0.4 ml/min. The data were acquired with Agilent MassHunter Workstation Software, LC/MS Data Acquisition B.08.00 (Build 8.00.8058.0) operating in Auto MS/MS mode whereby the 20 most intense ions (charge states, 2 to 5) within 300 to 1,700 m/z mass range above a threshold of 1,000 counts were selected for MS/MS analysis. MS/MS spectra (50 to 1,700 m/z) were collected with the quadrupole set to “narrow” resolution and were acquired until 25,000 total counts were collected or for a maximum accumulation time of 333 ms. To assure the desired mass accuracy of recorded ions, continuous internal calibration was performed during analyses with the use of signals m/z 322.0481 (m/z 322.0481 (detected m/z [C₆H₁₈N₃O₆P₃ − H]⁺) and 1221.9906 (detected m/z [C₂₄H₁₈O₆N₃P₃F₃₆ − H]⁺)

Each MS/MS spectra data set was processed to determine monoisotopic masses and charge states, to merge MS/MS spectra with the same precursor (Δm/z < 1.4 Da and chromatographic Δt < 15 s) and to select high-quality spectra with the extraction tool of SpectrumMill Proteomics Workbench (Agilent Technologies, CA, USA). The reduced data set was searched against the *V. mediterranei* proteins database in the identity mode with the MS/MS search tool of SpectrumMillProteomics Workbench and with the following settings: trypsin, up to 2 missed cleavages, carbamidomethylation of cysteines as fixed modifications, oxidation of methionine as variable, mass tolerance of ±20 ppm for precursor and ±50 ppm for product ions. The precursor mass shift was set between -18 Da to 177 Da to take into consideration variable modifications such as the presence of sodium and potassium adducts. Peptide hits were validated in the peptide mode to achieve a false discovery rate (FDR) of <1.2% and then in the protein mode according to the score settings recommended by the manufacturer. Positive identifications were considered only when two or more peptides were matched, and their summed score was >20. For label-free relative quantitation, differentially expressed proteins were assessed based on the regulation of the peptides. Protein lists of all samples were exported to the Mass Profiler Professional (MPP) software for statistical data analysis. Data analysis was carried out based on the total spectra intensity of the proteins which were considered as entities in MPP. The baseline of the spectra was adjusted to the median across all samples. The entities were then filtered based on their frequency of occurrence across at least all replicates of one treatment. Two-way ANOVA test was performed with the P<0.05. Benjamini-Hochberg corrected P-values were calculated to overcome the problem of multiple test analysis (false discovery). Regulated proteins with ≥2.0-fold change were then considered as regulated proteins.

### Experiment with corals

Fragments (∼5 cm in diameter) of the coral *O. patagonica* were collected in December 2024 from Alicante Harbour (38°20′11.1″ N, 0°29′11.8″ W) and transferred immediately to the laboratory. Coral fragments were randomly assigned to one of six aquaria (20 L each), with six pseudoreplicated fragments per aquarium, containing sterile seawater, and acclimated for 3 days under laboratory conditions. Aquaria were equipped with daylight LED lamps (4.8 W, 9000 K), programmed to follow the natural day–night cycle at a constant light intensity corresponding to that time of year, and aeration was provided using air pumps. Following acclimation, coral fragments were gradually adjusted to the experimental temperature by increasing seawater temperature by 0.5 °C per day, from 22 °C (the seawater temperature at the sampling site) to 28 °C. Once the target temperature was reached, corals were maintained at 28 °C for an additional 3 days prior to inoculation. Six different treatments were tested, consisting of non-inoculated corals (control), corals infected with *V. mediterranei*, corals maintained with 0.1 mM DMSP, corals maintained with 0.1 mM DMSP and infected with *V. mediterranei*, corals infected with *V. mediterranei* and *P. adelos*, and corals maintained with 0.1 mM DMSP and infected with both *V. mediterranei* and *P. adelos*. In all treatments involving inoculation, *V. mediterranei* and *P. adelos* were added at final concentrations of 10³ CFUs ml⁻¹ and 10² PFU ml⁻¹, respectively (i.e., MOI=0.1, as in the one-step experiment).

After 17 d of incubation, coral fragments were gently washed three times with 50 ml of sterile- filtered seawater (SFSW) to remove non-associated bacteria. Coral fragments were then broken into small pieces, placed into 50 ml tubes, and centrifuged for 3 min at 2,900×g to separate mucus from coral tissue and skeleton. Coral mucus was collected and used for *Vibrio* and phage counts, as well as for DNA extraction. For plate counts of *Vibrio* spp., 10-fold serial dilutions of coral mucus were prepared in SFSW, plated onto TCBS agar (Pronadisa, Spain), and incubated at 30 °C for 48 h. Enumeration of phage infecting *V. mediterranei* strain Vic-Oc-097 was carried out by plaque assay as described above.

After mucus extraction, the remaining coral pieces were crushed in SFSW using a mortar and pestle. The CaCO₃ skeleton was allowed to settle for 15 min, after which the supernatant (coral tissue homogenate) was collected for chlorophyll *a* (Chl *a*) determination. For Chl *a* measurements, 1 ml of the homogenate was centrifuged at 5,000×g for 10 min at 4 °C, and the supernatant was discarded. Pigments were extracted from the resulting pellet by incubation in 10 ml of 90% acetone at 4 °C for 24 h in the dark, followed by centrifugation at 13,000×g for 10 min. Absorbance was measured at 750, 664, 647, and 630 nm, and Chl *a* concentrations were calculated following Jeffrey and Humphrey (1975).

### DNA extractions and sequencing

Prior to DNA extraction, 350 µl of mucus obtained from coral colonies were incubated with lysozyme (0.4 mg/ml) for 30 min at 37 °C. DNA was extracted with the DNeasy Blood & Tissue Kit (Qiagen, Germany) following the manufacturer’s protocol and eluted in 35 μl of ultrapure water. DNA was quantified using a Qubit 2.0 Flourometer (Life Technologies, California, USA. Extracted DNA was stored at -80 °C until use.

The extracted DNA was used for PCR amplification of the V3–V4 regions of the 16S rRNA gene using primers 341F–805R and 25 amplification cycles (Takahashi et al., 2014). PCR reactions were prepared in a final volume of 50 µl containing: 1.5 mM MgCl₂, 10× buffer, 10 mM dNTPs, 5 U/μl Taq polymerase, 100 μM primers, 5 ng of the template DNA and ultrapure water up to the volume. Five ml of the amplified products were electrophoresed, stained with ethidium bromide (100 μg/ml), and visualized under UV light. The remaining products were sequenced on an Illumina Miseq (2 x 150 bp) at Fundació per al Foment de la Investigació Sanitària i Biomèdica de la Comunitat Valenciana (Fisabio, Valencia, Spain).

### Metabarcoding and statistical analyses

Illumina sequences were processed following the QIIME2 pipeline (Bolyen et al., 2019). Briefly, after trimming all reads using the cutadapt integrated tool in QIIME2, low-quality sequences were removed (sequences with less than 33 in Phred quality score). Sequences were clustered into Operational Taxonomic Units (OTUs) that were defined at the 98.7% identity, threshold previously used to distinguish species (Bolyen et al. 2019). Taxonomy assignments were carried out using the SILVA reference database (version 138) performing the machine learning algorithm Scikit-learn in QIIME2 (Stackebrandt and Ebers 2006). Chimera sequences were removed, and a frequency table was obtained using vsearch algorithm (Pedregosa et al. 2011) implemented in QIIME2.

Amplicon sequence data were analyzed using compositional data analysis (CoDA) approaches(Gloor et al., 2017), following the same analytical framework previously applied by our group (Rubio-Portillo et al., 2026). Prior to analysis, low-abundance OTUs were filtered to reduce noise, and zero counts were addressed using Bayesian-multiplicative replacement as implemented in the *zCompositions* R package. All analyses were conducted in R (version 4.4.3). To identify OTUs associated with differences between experimental conditions, three complementary CoDA-based methods were applied: generalized linear models with elastic net regularization (*glmnet*) (Friedman et al., 2010), log-ratio analysis (LogRatios) implemented in the *coda4microbiome* package (Calle et al., 2023), and ALDEx2, which applies centered log-ratio (CLR) transformation combined with Monte Carlo sampling from the Dirichlet distribution to estimate differential abundance while accounting for compositional constraints (Fernandes et al., 2014). OTUs identified by either method were retained for downstream interpretation.

### Statistical analysis and data visualization

Statistical analyses and data visualization were performed in R (v. 4.4.3) using the *dplyr*, *tidyr*, *ggplot2*, *ggpubr*, *scales*, *combinat*, *glmnet*, *coda4microbiome*, *ALDEx2* and *zCompositions* packages, as applicable. Unless otherwise stated, data were presented as individual observations together with the mean ± standard error of the mean (SEM). For AFM particle-volume data, individual detected particles are shown together with box plots representing the median and interquartile range, with whiskers extending to 1.5 × the interquartile range. Missing values were excluded on an analysis-by-analysis basis. All statistical tests were two-sided, and P < 0.05 was considered statistically significant. Exact P values were reported where applicable.

Comparisons between two independent groups were performed using two-sided unpaired Welch’s t-tests. For the phage infectivity assay, each DMSP concentration was compared with the untreated control separately for each phage, and P values were adjusted by the Bonferroni method for the four comparisons. AFM particle volumes in control and DMSP-treated samples were compared using a two-sided unpaired Welch’s t-test. For bacterial growth experiments, control and DMSP- treated cultures were compared separately at the indicated sampling times using two-sided unpaired Welch’s t-tests. Exact P values are shown for the prespecified late sampling points, including values above the significance threshold, to indicate the magnitude of the observed differences. In the coral experiment, phage abundance in mucus was compared between the two treatments containing both *V. mediterranei* and *P. adelos*, with or without DMSP, using a two- sided unpaired Welch’s t-test. Chlorophyll *a* content and Shannon diversity were assessed using all pairwise two-sided unpaired Welch’s t-tests among treatments, followed by Bonferroni correction for multiple comparisons. Only adjusted P values were used to determine statistical significance.

Microbiome differential-abundance analyses were performed using the CoDA framework described above. Differences in coral microbial community composition were assessed using permutational multivariate analysis of variance (PERMANOVA; Anderson, 2001), implemented in the adonis2 function of the R package *vegan*, with Vibrio, DMSP and Phage as explanatory factors. Aitchison distances were calculated as Euclidean distances between centred log-ratio (CLR)-transformed community profiles following zero replacement. Model terms were tested sequentially using 9999 unrestricted permutations. Homogeneity of multivariate dispersions was assessed using betadisper and the corresponding permutation test. Statistical significance was defined as p < 0.05.

## ACKNOWLEDGEMENTS

This research was supported by the project CONNECTIVITY PID2024-158829NB-C41 to F.S. and J.A. (Spanish Ministry of Science and Innovation). R.S.M. received funding for his doctoral thesis from the Spanish Ministry of Science and Innovation PRE2019-087998. R.S.M., E.R.P., F.S. and J.A. are members of the National Excellence Network FAGOMA (RED2022-134837-T).

## ETHICS DECLARATIONS

The authors declare no competing interests.

---

**Extended Data Fig. 1.**
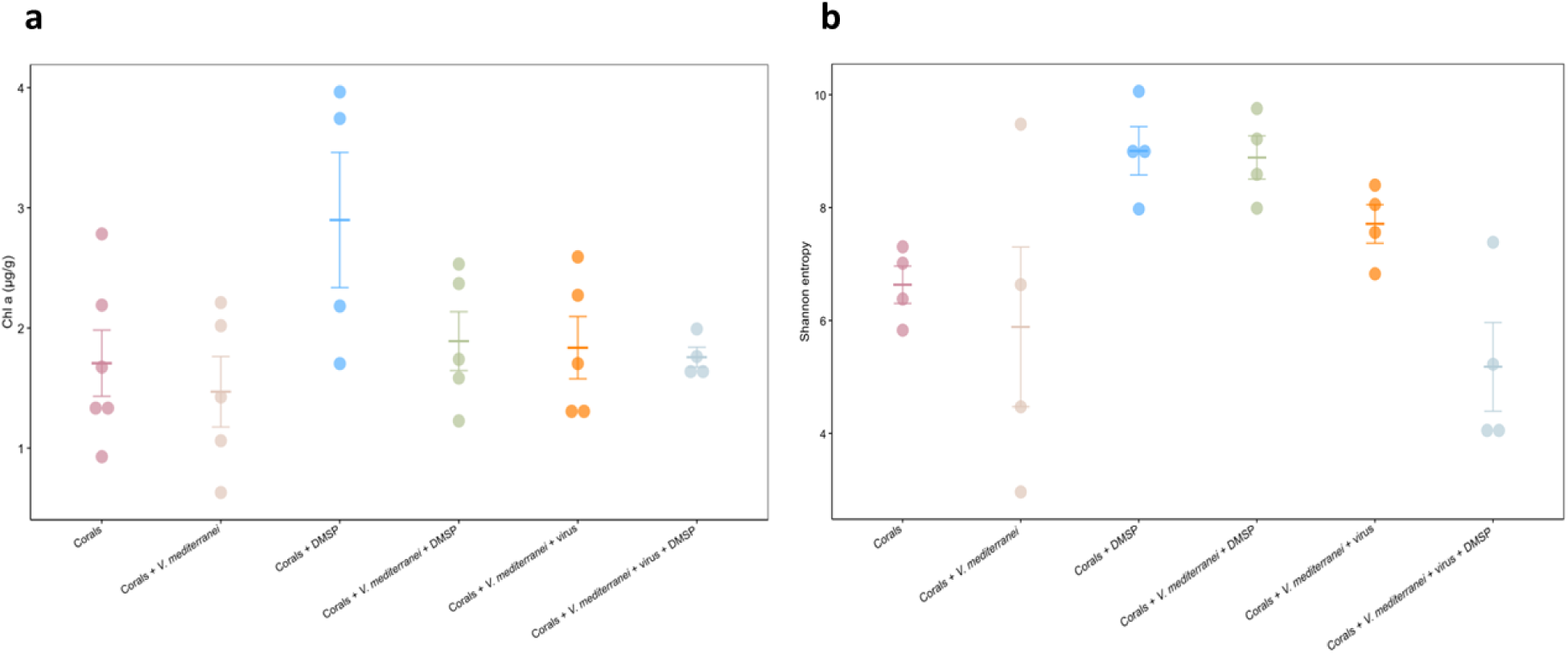
: Physiological status and alpha diversity of coral fragments subjected to the aquarium treatments. a, Chlorophyll *a* content of *O. patagonica* fragments after 17 days under the six experimental treatments. **b,** Shannon diversity of the coral mucus-associated prokaryotic communities determined by 16S rRNA gene metabarcoding. Dots represent individual coral fragments; horizontal lines and error bars indicate the mean ± SEM. All pairwise comparisons between treatments were performed using two-sided unpaired Welch’s t-tests, followed by Bonferroni correction for multiple comparisons. No statistically significant differences were detected after correction (adjusted P > 0.05).

## Notes

### Competing Interest Statement

The authors have declared no competing interest.

