## Supplementary Material for "DMSP modulates virus-host interactions in the coral environment"

July 21, 2026

These notes present in a constructive way the development of a formal dynamical model tailored to the empirical observations reported in the main ms (three first subsections in RESULTS section). We begin by introducing a minimal model (S1) initially motivated by the observed burst in viral production with and withouth DMSP. Subsequently, we analytically derive conditions for reproducing the reversal observed at larger times, and show how the simplest DMSP hypothesis falls short (S2). Then, parameters of an extended model (S3) are derived to fit to one-step-curve data (S4).

#### S1 A minimal model of infection dynamics

We propose a system of three differential equations modeling the dynamics of susceptible $S(t)$ and infected $I(t)$ cells, and the amount of free virions $V(t)$. The model constitutes a particular case of the general Levin–Stewart–Chao (LSC) model to simulate how bacterial populations grow, compete for limited nutrients, and interact with predatory bacteriophages [1]. The model has five parameters: the duplication rate $r$ of susceptible cells, the infection (or adsorption) rate $\phi$, the burst size $B$, the latent period (lysis time) $\tau$, and the virion decay rate $\omega$:

$$\dot{S} = rS(t) - \phi V(t)S(t), \tag{1}$$

$$\dot{V} = -\phi V(t)S(t) + B\,\phi V(t-\tau)S(t-\tau) - \omega V(t), \tag{2}$$

$$\dot{I} = \phi V(t)S(t) - \phi V(t-\tau)S(t-\tau). \tag{3}$$

With respect to the original LSC model, we assume:

1. A carrying capacity $K = \infty$. This is consistent with the experiments, since they last only a few hours and nutrients are not limiting in the relevant dynamical region;
2. Cell death is produced only by lysis, so the mortality of infected bacteria is set to $m = 0$, eliminating an exponential term in the LSC model;
3. Lysis time is fixed, $\tau = \text{ct}$. This mimics the step rise of one-step curves.

#### S2 First hypothesis: DMSP acts only through $\tau$ and $B$

An initial set of experiments suggested that DMSP could be acting on burst size and on lysis time: $B$ decreased and $\tau$ increased with respect to control experiments (Figure 3a). Simulations of Eqs. (1)–(3) implementing this hypothesis, that is, larger values of $\tau$ and smaller values of $B$ trivially reproduce a delay in the burst time and a lower burst size.

However, when experiments are extended for longer times, a reversal in the amount of viral particles produced is observed. That is, the long-time production of viral particles is higher in the system treated with DMSP, despite the lower value of $B$ and higher value of $\tau$ that reproduces the dynamics until $t \simeq 120$ minutes.

In the following of this section, we show through analytic and heuristic arguments that the observed reversal cannot occur under the hypothesis that DMSP only affects $\tau$ and $B$. The summary of the reasoning is as follows: we first calculate the basic reproduction number $R_0$ and show that it does not depend on the lysis time $\tau$, but does on the $B$ and on the number of susceptible cells $S$. Increasing $\tau$ therefore affects the number of susceptible cells available at lysis time; since this time is larger for the bacteria-phage system in presence of DMSP, the number of susceptible cells is also larger, potentially increasing the number of efficient infections even if the burst size is smaller. We perform a heuristic extension of the $R_0$ derivation to show that, at lysis time, viral growth rate in presence of DMSP could be larger than the control growth rate if the difference in lysis time between the two systems is large enough, so as to produce an increase in susceptible cells that could compensate for the disadvantage of a lower burst. The use of empirical data reveals that the duplication time of the bacteria had to be unrealistically high to cause the compensation. We therefore conclude that only if DMSP also affects the host can the reversal in virus production occur as observed.

##### S2.1 Local invasion analysis: basic reproduction number $R_0$

In general, the basic reproduction number $R_0$ of an infection establishes the likelihood that a viral invasion succeeds in an infection-free host population. This amounts to setting $V \simeq 0$ and $S \simeq S^*$, constant. In this way, the equation for the growth of the number of viral particles $\dot{V}$ becomes linear,

$$\dot{V} = -(\phi S^* + \omega)V(t) + B\,\phi S^* V(t-\tau)\,. \tag{4}$$

We seek a solution of the form $V \sim e^{\lambda t}$ and find $\lambda = -(\phi S^* + \omega) + B\phi S^* e^{-\lambda\tau}$, yielding

$$R_0 = \frac{B\phi S^*}{\phi S^* + \omega}\,. \tag{5}$$

That is, the critical threshold where the viral population changes from growing to declining, $\lambda = 0$, is equivalent to $R_0 = 1$. Since $\tau$ does not appear in the expression of $R_0$ and this quantity varies monotonically with $B$ and $S^*$, it cannot change the sign of $\lambda$ at fixed $S^*$.

##### S2.2 Heuristic extension of the threshold analysis

The threshold condition derived above allows comparison among systems evaluated at the same bacterial density $S^*$. Consequently, if two viral strains differ only in burst size, the strain with the larger burst size always has the larger basic reproduction number. Likewise, if both strains experience the same susceptible host density, increasing $S^*$ benefits both systems equally and cannot reverse their ordering.

However, this comparison does not correspond exactly to the biological situation considered in our work. The local threshold analysis assumes that the susceptible population size is fixed, whereas in the complete model the bacterial population continues growing during the latent period represented by the lysis time $\tau$. If two viral phenotypes have different latent periods, they therefore experience different effective host densities when progeny virions are released.

This observation motivates the following heuristic extension of the threshold analysis. In the following, the subindex $C$ stands for control systems and $D$ for systems in presence of DMSP.

Suppose that DMSP modifies only viral traits by decreasing the burst size, $B_D < B_C$ and increasing the latent period, $\tau_D > \tau_C$, while leaving cellular growth rate $r$, adsorption rate and virion decay unchanged, $\phi_D = \phi_C = \phi$, $\omega_D = \omega_C = \omega$. Also assume (in agreement with empirical observations and the model equations) that susceptible bacteria grow exponentially, $S(t) = S_0 e^{rt}$.

Under this approximation, the susceptible density available at the time of the first lysis event is $S_C = S_0 e^{r\tau_C}$ for the control system, and $S_D = S_0 e^{r\tau_D}$ for the DMSP-treated system. Therefore,

$$\frac{S_D}{S_C} = e^{r(\tau_D - \tau_C)} > 1. \tag{6}$$

Equation (5) was originally derived assuming a constant susceptible density. We now use it heuristically by replacing the constant $S^*$ with the effective susceptible density experienced at the time of lysis. This is no longer a rigorous local stability calculation, but rather an approximate way of incorporating the increase in host abundance that occurs during the latent period. Under this approximation,

$$R_{0,C} = \frac{B_C \phi S_C}{\phi S_C + \omega}, \qquad R_{0,D} = \frac{B_D \phi S_D}{\phi S_D + \omega}. \tag{7}$$

Taking the ratio,

$$\frac{R_{0,D}}{R_{0,C}} = \frac{B_D}{B_C} \frac{S_D}{S_C} \frac{\phi S_C + \omega}{\phi S_D + \omega}. \tag{8}$$

Equation (8) separates naturally into three factors: the reduction in burst size, $B_D/B_C$, the increase in susceptible host density, $S_D/S_C$, and a correction factor arising from the saturation of $R_0$ (last term on the rhs). Introducing $x = \phi S_C$ and $k = S_D/S_C > 1$, Eq. (8) becomes

$$\frac{R_{0,D}}{R_{0,C}} = \frac{B_D}{B_C} k \frac{x + \omega}{kx + \omega}. \tag{9}$$

Simple algebra now yields useful bounds. First, $kx + \omega > x + \omega$, which implies

$$\frac{R_{0,D}}{R_{0,C}} < \frac{B_D}{B_C} \frac{S_D}{S_C}. \tag{10}$$

Second, $kx + \omega < k(x + \omega)$, therefore

$$\frac{B_D}{B_C} < \frac{R_{0,D}}{R_{0,C}} < \frac{B_D}{B_C} \frac{S_D}{S_C}. \tag{11}$$

Equation (11) shows that bacterial growth during the additional latent period always increases the ratio $R_{0,D}/R_{0,C}$ above the burst-size ratio alone, but that the improvement is bounded above by the increase in host density.

A necessary condition for the DMSP-treated system to have a larger effective reproduction number than the control is $R_{0,D} > R_{0,C}$. Using the upper bound (10), this requires $\frac{B_D}{B_C}\frac{S_D}{S_C} > 1$ or, equivalently,

$$\frac{S_D}{S_C} > \frac{B_C}{B_D}. \tag{12}$$

Substituting Eq. (6) gives $e^{r(\tau_D - \tau_C)} > B_C/B_D$,which can also be written as

$$r(\tau_D - \tau_C) > \ln\left(\frac{B_C}{B_D}\right). \tag{13}$$

Equation (13) provides a simple biological interpretation. The increase in host availability produced during the additional latent period must exceed the reduction in burst size for the DMSP-treated system to overcome the intrinsic disadvantage associated with producing fewer progeny.

We emphasize that this result is not a rigorous property of the local threshold analysis. The quantity $R_0$ was originally derived assuming a constant susceptible density. Equations (8)–(13) therefore represent a heuristic extension in which the constant density entering the threshold expression is replaced by the effective density experienced at lysis. The argument should thus be regarded as an approximate way of quantifying the competition between two opposing effects of DMSP: the reduction in burst size and the increase in host availability resulting from a longer latent period.

##### S2.3 Application to the experimental measurements

The measured latent periods are $\tau_C = 79.9$ min and $\tau_D = 84.2$ min, so that $\Delta\tau = \tau_D - \tau_C = 4.3$ min. The measured burst-size ratio is $\frac{B_D}{B_C} = 0.78$, corresponding to a reduction of approximately 22% under DMSP.

Using Eq. (13), the necessary condition for the DMSP-treated system to attain a larger effective reproduction number becomes

$$e^{r\Delta\tau} > \frac{1}{0.78} = 1.282\,, \tag{14}$$

which yields $r > 0.0577$ min$^{-1}$. Expressed as a bacterial doubling time, $T_d = \frac{\ln 2}{r}$, we get

$$T_d < \frac{0.693}{0.0577} \simeq 12.0 \text{ min}\,. \tag{15}$$

This value is considerably shorter than typical doubling times reported for marine bacteria, shorter than the fastest growth rates commonly observed for laboratory cultures of *Escherichia coli* under optimal conditions, and certainly below the measured doubling time of 37 min for *V. mediterranei*[1]. For a $T_d = 37$ min, the growth rate $r = 0.0187$ min$^{-1}$, giving $e^{r\Delta\tau} = 1.084$, which is below the value required by Eq. (13).

Therefore, under the assumptions adopted in this heuristic extension—namely, that DMSP modifies only the burst size and latent period while cellular growth rate, adsorption and virion decay remain unchanged—the increase in susceptible host density produced during the additional 4.3 minutes of latency appears insufficient to compensate for the measured reduction in burst size unless bacterial growth is unrealistically fast, which is not the case. Although bacterial growth during the latent period acts in the appropriate direction, regarding reversal at later times, by increasing the effective host density, the measured changes in burst size and latent period indicate that this mechanism alone is unlikely to explain the experimentally observed long-term reversal. Consequently, additional DMSP-induced effects, such as changes in adsorption efficiency, virion survival, or other mechanisms that increase the effective infection efficiency, would appear necessary if the reversal is to be explained within the framework of the present model.

[1]Here we should say something about the origin of this value, whether it is unpublished (and the journal accepts it) or show data supporting that value.

### S3 Extended model: protocol structure and priming

Section S2 establishes that changes in $\tau$ and $B$ alone cannot plausibly account for the observed reversal: the compensation would require a bacterial doubling time under 12 minutes, well outside the physiological range. An additional mechanism affecting the effective infection efficiency – most simply, an increase in the adsorption rate $\phi$ itself – therefore appears necessary. Before fitting such a mechanism to data, we first refine the model to match the structure of the experimental protocol that generated the one-step-curve measurements, since Eqs. (1)–(3) as written assume continuous, ongoing infection from $t = 0$, which is not what the assay does.

The one-step-curve protocol infects host cells at a fixed multiplicity of infection (MOI, $m = 0.1$), allows 5 minutes for adsorption, and then dilutes the mixture $10^2$–$10^4$-fold, specifically to prevent further adsorption over the subsequent observation window (Methods). The dynamics therefore proceed in two phases. During **Phase 1** ($0 \leq t \leq \tau$), the single cohort of cells infected during the initial adsorption window ages synchronously and releases its progeny in one burst at $t = \tau$; no new infections occur, since free virions and uninfected cells alike have been diluted below the density at which further encounters are likely:

$$S(t) = S_0 e^{rt}, \qquad I(t) = mS_0, \qquad V(\tau^+) = BmS_0 \qquad (0 \leq t \leq \tau), \tag{16}$$

where $S_0$ is the initial concentration (or number) of susceptible cells (immediately after the experimentally aplied dilution).

**Phase 2** ($t > \tau$) begins once the culture is left to incubate further and re-infection resumes, seeded by the Phase-1 burst: $S(\tau^+) = S_0 e^{r\tau}$, $V(\tau^+)$ as in Eq. (16). Equations (1)–(3) apply from this point on, with one addition: since the mechanism motivated at the start of this section is an increase in adsorption efficiency, and since the proteomic data (main text) show the relevant receptor-abundance changes becoming detectable only 150–180 minutes after DMSP exposure, we let $\phi$ depend on time in the DMSP-treated system,

$$\phi(t) = \phi_{\text{naive}} + \left(\phi_{\text{primed}} - \phi_{\text{naive}}\right) \Big/ \left(1 + e^{-(t-t_c)/w}\right), \tag{17}$$

a smooth (logistic) switch centered at $t_c \approx 165$ min, the midpoint of the reported 150–180 min window, with width $w = 15$ min. In the control system $\phi(t) = \phi_{\text{naive}}$ throughout. Equation (17) is a population-averaged description: it represents the mean adsorption efficiency of the culture as a whole gaining sensitivity over time, rather than an explicit split between a "naive" and a "primed" subpopulation of cells, which the bulk proteomic measurements available do not resolve.

### S4 Fit to empirical data

#### S4.1 Data

Triplicate measurements were provided for a fine-resolution one-step curve (5-minute steps, $t = 5$–110 min, used to determine $\tau$ and the burst-jump ratio) and a long-term series ($t = 0$–1510 min, used to determine the secondary-phase growth rate and the crossover time). Table S1 and Table S2 report the triplicate mean ± standard deviation at every measured timepoint. We note that the two sets of measurements are not on a common absolute scale (their ratio at the three shared timepoints, $t = 65, 85, 105$ min, is not constant), consistent with these being independent experimental runs; all quantities extracted below are therefore scale-invariant (ratios, rates, timings), never raw magnitudes compared across experiments.

Table S1: One-step curve, 5-min resolution (sheet "Burst size"). PFU in raw (undiluted-equivalent) assay units; mean ± SD, $n = 3$.

| $t$ (min) | Control | DMSP |
|---|---|---|
| 5 | $1.88{\times}10^3 \pm 5.2{\times}10^2$ | $1.44{\times}10^3 \pm 4.4{\times}10^2$ |
| 10 | $2.02{\times}10^3 \pm 5.8{\times}10^2$ | $1.52{\times}10^3 \pm 4.3{\times}10^2$ |
| 15 | $1.96{\times}10^3 \pm 4.7{\times}10^2$ | $1.45{\times}10^3 \pm 4.6{\times}10^2$ |
| 20 | $2.09{\times}10^3 \pm 5.0{\times}10^2$ | $1.13{\times}10^3 \pm 1.0{\times}10^3$ |
| 25 | $2.03{\times}10^3 \pm 6.5{\times}10^2$ | $1.63{\times}10^3 \pm 3.5{\times}10^2$ |
| 30 | $1.97{\times}10^3 \pm 6.1{\times}10^2$ | $1.67{\times}10^3 \pm 4.7{\times}10^2$ |
| 35 | $2.87{\times}10^3 \pm 5.5{\times}10^2$ | $1.87{\times}10^3 \pm 8.1{\times}10^2$ |
| 40 | $3.00{\times}10^3 \pm 7.2{\times}10^2$ | $2.27{\times}10^3 \pm 7.8{\times}10^2$ |
| 45 | $3.37{\times}10^3 \pm 9.1{\times}10^2$ | $2.63{\times}10^3 \pm 7.6{\times}10^2$ |
| 50 | $3.40{\times}10^3 \pm 1.2{\times}10^3$ | $3.13{\times}10^3 \pm 6.0{\times}10^2$ |
| 55 | $3.00{\times}10^3 \pm 1.0{\times}10^3$ | $2.67{\times}10^3 \pm 5.8{\times}10^2$ |
| 60 | $3.33{\times}10^3 \pm 5.8{\times}10^2$ | $3.67{\times}10^3 \pm 5.8{\times}10^2$ |
| 65 | $8.33{\times}10^3 \pm 1.5{\times}10^3$ | $7.33{\times}10^3 \pm 5.8{\times}10^2$ |
| 70 | $4.93{\times}10^4 \pm 8.5{\times}10^3$ | $9.00{\times}10^3 \pm 1.0{\times}10^3$ |
| 75 | $3.77{\times}10^5 \pm 6.5{\times}10^4$ | $1.03{\times}10^4 \pm 4.2{\times}10^3$ |
| 80 | $1.30{\times}10^6 \pm 1.2{\times}10^5$ | $6.60{\times}10^5 \pm 1.1{\times}10^5$ |
| 85 | $1.70{\times}10^6 \pm 1.9{\times}10^5$ | $1.05{\times}10^6 \pm 8.2{\times}10^4$ |
| 90 | $2.44{\times}10^6 \pm 1.4{\times}10^5$ | $1.40{\times}10^6 \pm 1.5{\times}10^5$ |
| 95 | $2.52{\times}10^6 \pm 8.7{\times}10^4$ | $1.86{\times}10^6 \pm 1.3{\times}10^5$ |
| 100 | $2.54{\times}10^6 \pm 6.4{\times}10^4$ | $2.01{\times}10^6 \pm 5.1{\times}10^4$ |
| 105 | $2.58{\times}10^6 \pm 5.9{\times}10^4$ | $2.02{\times}10^6 \pm 8.5{\times}10^4$ |
| 110 | $2.53{\times}10^6 \pm 4.0{\times}10^4$ | $2.01{\times}10^6 \pm 1.0{\times}10^5$ |

Table S2: Long-term series (sheet "Burst size long"), PFU/mL, mean ± SD, $n = 3$.

| $t$ (min) | Control | DMSP |
|---|---|---|
| 0 | $6.36{\times}10^5 \pm 3.1{\times}10^4$ | $6.25{\times}10^5 \pm 2.7{\times}10^4$ |
| 15 | $6.94{\times}10^5 \pm 4.7{\times}10^4$ | $6.79{\times}10^5 \pm 2.0{\times}10^4$ |
| 30 | $6.03{\times}10^5 \pm 1.5{\times}10^4$ | $6.27{\times}10^5 \pm 6.1{\times}10^4$ |
| 65 | $1.47{\times}10^6 \pm 1.1{\times}10^5$ | $7.20{\times}10^5 \pm 4.0{\times}10^4$ |
| 85 | $2.03{\times}10^7 \pm 1.4{\times}10^6$ | $3.40{\times}10^6 \pm 2.0{\times}10^5$ |
| 105 | $2.42{\times}10^8 \pm 3.6{\times}10^7$ | $1.44{\times}10^8 \pm 8.3{\times}10^6$ |
| 120 | $2.51{\times}10^8 \pm 1.9{\times}10^7$ | $1.95{\times}10^8 \pm 6.0{\times}10^6$ |
| 155 | $3.67{\times}10^8 \pm 4.9{\times}10^7$ | $5.00{\times}10^8 \pm 7.0{\times}10^7$ |
| 180 | $3.50{\times}10^8 \pm 6.2{\times}10^7$ | $9.60{\times}10^8 \pm 2.6{\times}10^7$ |
| 240 | $6.73{\times}10^8 \pm 7.5{\times}10^7$ | $1.12{\times}10^9 \pm 4.4{\times}10^7$ |
| 300 | $9.17{\times}10^8 \pm 9.3{\times}10^7$ | $2.37{\times}10^9 \pm 7.5{\times}10^7$ |
| 330 | $2.03{\times}10^9 \pm 3.1{\times}10^8$ | $4.40{\times}10^9 \pm 2.0{\times}10^8$ |
| 400 | $3.87{\times}10^9 \pm 6.5{\times}10^8$ | $9.47{\times}10^9 \pm 9.3{\times}10^8$ |
| 430 | $7.13{\times}10^9 \pm 6.5{\times}10^8$ | $1.34{\times}10^{10} \pm 1.2{\times}10^9$ |
| 460 | $8.00{\times}10^9 \pm 4.0{\times}10^8$ | $1.84{\times}10^{10} \pm 1.2{\times}10^9$ |
| 500 | $9.67{\times}10^9 \pm 1.3{\times}10^9$ | $2.72{\times}10^{10} \pm 1.9{\times}10^9$ |
| 530 | $1.30{\times}10^{10} \pm 1.8{\times}10^9$ | $3.18{\times}10^{10} \pm 3.4{\times}10^9$ |
| 1250 | $1.63{\times}10^{11} \pm 3.1{\times}10^{10}$ | $3.10{\times}10^{11} \pm 6.0{\times}10^{10}$ |
| 1330 | $2.43{\times}10^{11} \pm 4.0{\times}10^{10}$ | $2.90{\times}10^{11} \pm 2.6{\times}10^{10}$ |
| 1510 | $2.60{\times}10^{11} \pm 2.6{\times}10^{10}$ | $3.03{\times}10^{11} \pm 2.3{\times}10^{10}$ |

##### S4.2 Parameter values extracted from the data

We already extracted $\tau_C = 79.9$ min and $\tau_D = 84.2$ min as the 50%-rise time between the pre-burst baseline ($t$ = 5–60 min) and the post-burst plateau ($t$ = 95–110 min) in Table S1 (closely matching the ∼80 min reported in the main text). The burst-jump ratio (plateau minus baseline, DMSP/control) is $1.97 \times 10^6/2.54 \times 10^6 = 0.776$, matching the raw burst-size ratio $356/455 = 0.782$ closely.

From Table S2, a log-linear fit over $t$ = 105–530 min (the multi-cycle, pre-plateau regime) gives the secondary-phase growth rates $\lambda_C = 0.01000$ min$^{-1}$ and $\lambda_D = 0.01237$ min$^{-1}$, a fold-increase of 1.24. Interpolating the DMSP/control ratio to 1 (that is, finding the time when the two curves fitted to data cross each other) gives a crossover time somewhere in the 130–136 min range, bracketed by measurements at 120 and 155 min, consistent with Results' qualitative 'after 150 minutes.' At the latest measured timepoints ($t$ = 1250–1510 min), the DMSP/control ratio – which peaks at ∼2.1–2.8× around $t$ = 300–530 min – narrows to 1.17–1.9×: both cultures appear to be converging toward a shared ceiling at very late times rather than diverging indefinitely, plausibly reflecting exhaustion of the susceptible-cell pool in both conditions.

##### S4.3 The implied change in adsorption efficiency

Solving the characteristic equation (Eq. 5) for the $\phi S^*$ product implied by each fitted rate $\lambda$ (using the measured $B_D = 356$ directly) gives

$$\frac{\phi_{\text{primed}}}{\phi_{\text{naive}}} = 2.00. \tag{18}$$

This closely matches the manuscript's proteomics significance threshold ($\geq$ 2-fold change) – an independent, non-trivial cross-check between the dynamical fit and the proteomics criterion used to call candidate receptor proteins as regulated.

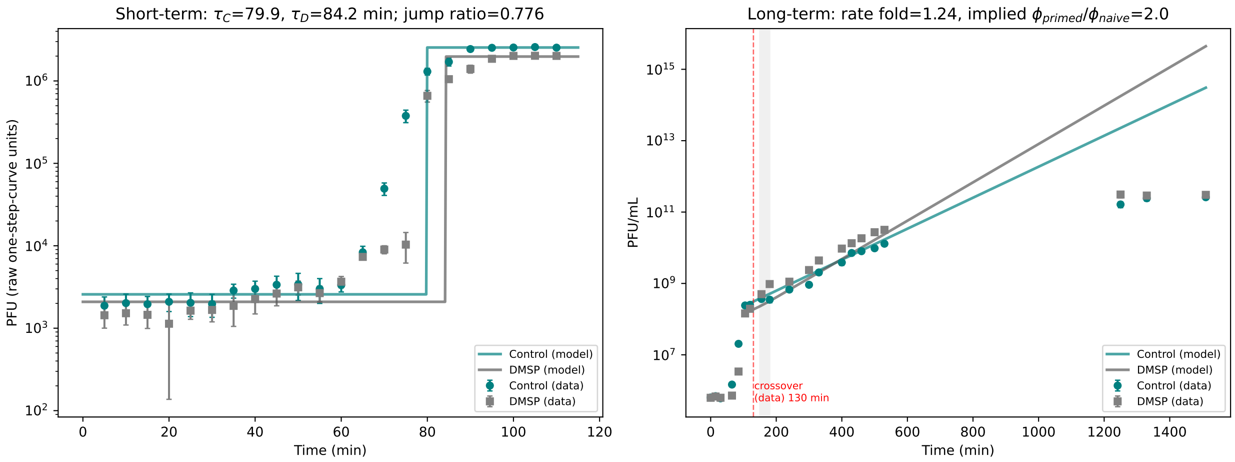


Figure S1: Model fit to the real triplicate one-step-curve data (Tables S1–S2). **Left panel** (short-term, $t = 0$–115 min): circles and squares are the measured mean $\pm$ SD ($n = 3$) for control and DMSP, respectively, in raw one-step-curve units (Table S1). The solid lines are the Phase-1 reconstruction of Eq. (16): flat at the pre-burst baseline until $t = \tau$ (79.9 min control, 84.2 min DMSP, vertical step), then flat at the post-burst plateau – by construction, since $\tau_C, \tau_D$ and the burst-jump ratio (0.776) are read directly from these same data. This panel visualizes the timing and relative size of the two bursts, not an independent prediction. **Right panel** (long-term, $t = 0$–1510 min, log-scale $y$-axis): circles and squares are the measured mean $\pm$ SD ($n = 3$) in PFU/mL (Table S2). Solid lines are the Phase-2 reconstruction: pure exponential growth at rate $\lambda_C$ (control) from the $t = 105$ min anchor, and exponential growth that switches smoothly (Eq. (17), shaded band = 150–180 min priming window) from $\lambda_C$ to $\lambda_D$ (DMSP) at $t_c = 165$ min. The vertical red dashed line marks the measured crossover time (≈130 min). This reconstruction is a direct fit of the two rates $\lambda_C, \lambda_D$ to the $t = 105$–530 min data (see main text of this section); it is not expected to, and does not, capture the late-time ($t > 1000$ min) narrowing of the DMSP/control ratio visible in the data, since it has no mechanism (such as susceptible-cell exhaustion) to bound growth.
